# Inhibition of Severe Acute Respiratory Syndrome Coronavirus 2 main protease by tafenoquine *in vitro*

**DOI:** 10.1101/2020.08.14.250258

**Authors:** Yeh Chen, Wen-Hao Yang, Li-Min Huang, Yu-Chuan Wang, Chia-Shin Yang, Yi-Liang Liu, Mei-Hui Hou, Chia-Ling Tsai, Yi-Zhen Chou, Bao-Yue Huang, Chian-Fang Hung, Yu-Lin Hung, Jin-Shing Chen, Yu-Ping Chiang, Der-Yang Cho, Long-Bin Jeng, Chang-Hai Tsai, Mien-Chie Hung

## Abstract

The severe acute respiratory syndrome coronavirus 2 (SARS-CoV-2) causing the current pandemic, coronavirus disease 2019 (COVID-19), has taken a huge toll on human lives and the global economy. Therefore, effective treatments against this disease are urgently needed. Here, we established a fluorescence resonance energy transfer (FRET)-based high-throughput screening platform to screen compound libraries to identify drugs targeting the SARS-CoV-2 main protease (M^pro^), in particular those which are FDA-approved, to be used immediately to treat patients with COVID-19. Mpro has been shown to be one of the most important drug targets among SARS-related coronaviruses as impairment of M^pro^ blocks processing of viral polyproteins which halts viral replication in host cells. Our findings indicate that the anti-malarial drug tafenoquine (TFQ) induces significant conformational change in SARS-CoV-2 M^pro^ and diminishes its protease activity. Specifically, TFQ reduces the α-helical content of M^pro^, which converts it into an inactive form. Moreover, TFQ greatly inhibits SARS-CoV-2 infection in cell culture system. Hence, the current study provides a mechanistic insight into the mode of action of TFQ against SARS-CoV-2 M^pro^. Moreover, the low clinical toxicity of TFQ and its strong antiviral activity against SARS-CoV-2 should warrant further testing in clinical trials.

## Introduction

The SARS-CoV-2 was identified in December 2019 as the cause of COVID-19 outbreak that originated in Wuhan, China (1–3). It has since spread rapidly, infected more than twelve million people globally, and caused more than 548,822 deaths (4). Currently, there are no scientifically proven drugs to control this outbreak. The SARS-CoV-2 genome shares about 83% identity with the SARS coronavirus that emerged in 2002 and contains approximately 30,000 nucleotides that are transcribed into 14 open reading frames (Orfs) (5). Among them, Orf1a and Orf1ab are translated into two polyproteins which are then cleaved by the main protease (M^pro^), yielding a number of protein products required for viral replication and transcription (6–9). Because there are no similar proteases in humans and that M^pro^ is necessary for viral replication, M^pro^ is considered as an ideal target for drug design. Drugs that are approved by the U.S. Food and Drug Administration (FDA) undergo rigorous evaluation for quality, safety and effectiveness, and thus identifying FDA-approved drugs that can inhibit M^pro^ protease activity has the advantage to be used quickly for treatment in patients. To this end, we established a fluorescence resonance energy transfer (FRET)-based high-throughput drug screening platform to rapidly identify antiviral compounds from the FDA-approved drug library which can bind to M^pro^ and inhibit its enzymatic activity.

## Results

### FRET-based high-throughput screening of FDA-approved compound library for inhibitors of SARS-CoV-2 M^pro^

Recombinant SARS-CoV-2 M^pro^ (ORF1ab polyprotein residues 3264–3569, GenBank code: QHD43415.1) was expressed in *Escherichia coli* and purified to homogeneity (fig. S1). To rapidly identify potential FDA-approved drugs targeting SARS-CoV-2 M^pro^, we established a fluorescence resonance energy transfer (FRET) assay by using a protein substrate consisting of the nsp4-5 N-terminal autocleavage site (TSAVLQ↓SGFRKM) of SARS-CoV-2 M^pro^ inserted between mTurquoise2 and mVenus, an enhanced CFP-YFP pair with higher quantum yield and protein stability (Fig. 1A and fig. S2) (10,11). The decrease in the FRET efficiency following cleavage of the protein substrate by SARS-CoV-2 M^pro^ and the increase in time-dependent fluorescence emission of mTurquoise2 at 474 nm were used as a measurement of M^pro^ activity. An initial screening of about 2,000 compounds using this FRET assay showed that tafenoquine (TFQ) exhibited the most significant inhibitory effect against SARS-CoV-2 M^pro^ (Fig. 1B).

**Fig. 1.**
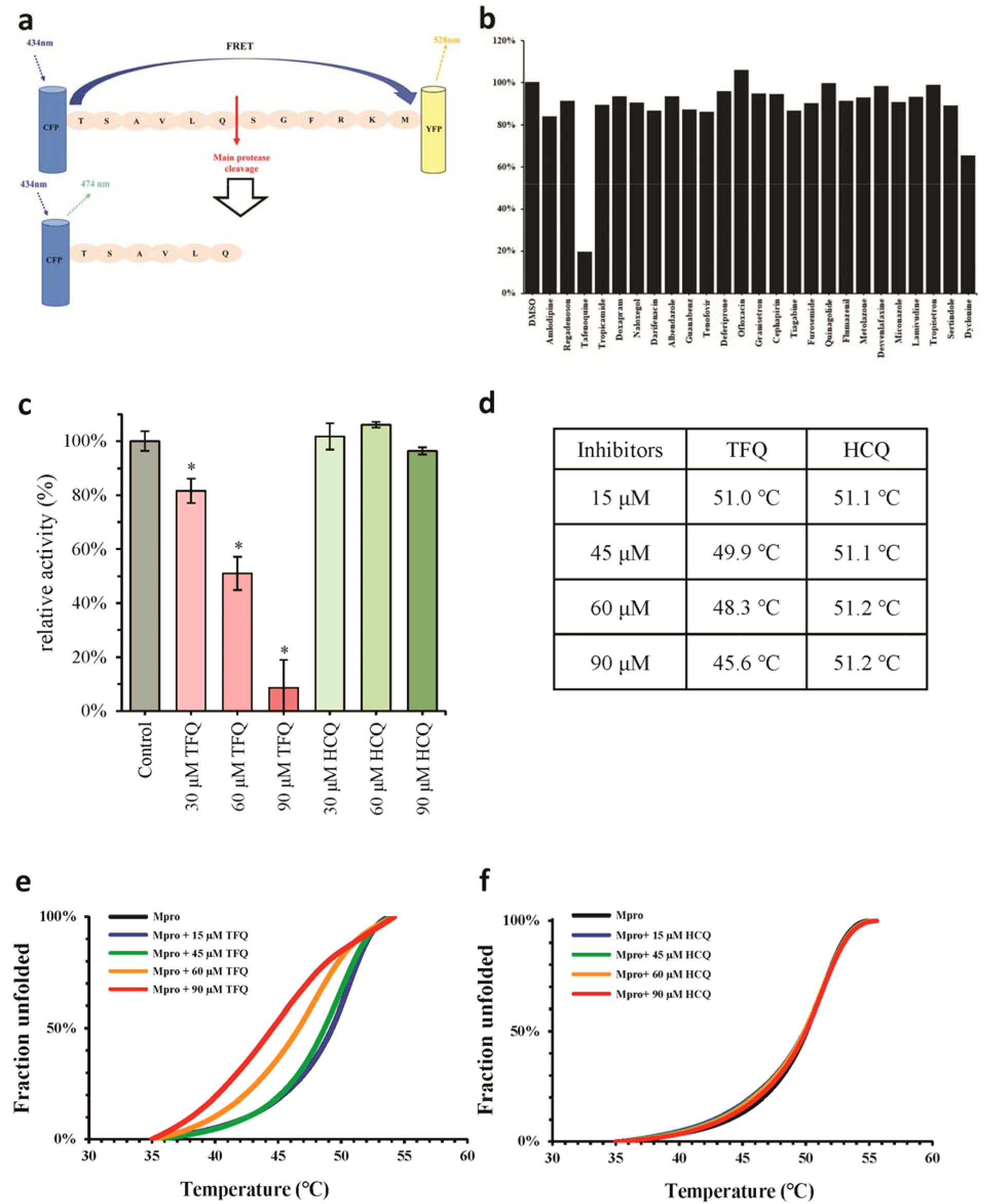
Drug repurposing screening of FDA-approved compound libraries against SARS-CoV-2 M^pro^. (**A**) Schematic presentation of FRET-based enzyme activity assay of SARS-CoV-2 M^pro^. (**B**) Drug repurposing screening of FDA-approved compound libraries against SARS-CoV-2 M^pro^ using FRET assay. Compounds (60 μM) were pre-incubated with 4 μM of SARS-CoV-2 M^pro^ for 30 min at room temperature. Substrates (20 μM) were then added to initiate the reaction. The relative enzyme activity of SARS-CoV-2 M^pro^ with 25 selected FDA-approved drugs are shown. (**C**) A comparison of the relative enzyme activity of SARS-CoV-2 M^pro^ at various concentrations of TFQ or HCQ (30, 60, 90 μM). (**D**) A comparison of the melting temperature (Tm) of SARS-CoV-2 M^pro^ at various concentrations of TFQ or HCQ (15, 45, 60, 90 μM). N.D., not detected. n = 2. Data are shown as mean ± standard deviation (SD). \**P* < 0.05 in Student *t* test. (**E**) Melting curves of SARS-CoV-2 M^pro^ at various concentrations of TFQ (15, 45, 60, 90 μM). (f) Melting curves of SARS-CoV-2 M^pro^ at various concentration of HCQ (15, 45, 60, 90 μM).

### Mechanism of action of TFQ against SARS-CoV-2 M^pro^

TFQ (brand name Krintafel/Kozenis in U.S./Australia, owned and developed by GlaxoSmithKline) is an 8-aminoquinoline anti-malarial drug that was approved by the U.S. FDA in July 2018 and the Australian Therapeutic Goods Administration (TGA) in September 2018 for the radical cure of *Plasmodium vivax* (12–14), a parasite that causes malaria. In addition, TFQ (brand name Arakoda/Kodatef in U.S./Australia, owned by 60 Degrees Pharmaceuticals) was later approved by the FDA and the TGA for malaria prophylaxis (14,15). However, the molecular target of TFQ is still unknown. Recently, two 4-aminoquinoline derivatives, chloroquine (CQ) and hydroxychloroquine (HCQ), were shown to be effective in inhibiting SARS-CoV-2 infection *in vitro* (16,17). Many clinical trials using CQ or HCQ to treat patients with COVID-19 have also been reported, but some have found no benefit and possible harm in patients (18,19). CQ is thought to inhibit virus entry by modifying glycosylation of ACE2 receptor and spike protein or by interfering with the pH-dependent endocytic pathway (20,21).

To further characterize TFQ, we compared the inhibitory effects of TFQ and HCQ against SARS-CoV-2 M^pro^ at various concentrations by FRET and differential scanning fluorimetry (DSF) (22). As shown in Figure 1c, TFQ exhibited almost 90% inhibition against SARS-CoV-2 M^pro^ at a concentration of 90 μM whereas HCQ did not demonstrate any significant inhibitory effects. Using a protein thermal shift assay, we showed that TFQ caused a negative shift in the melting temperature (Tm) of SARS-CoV-2 M^pro^ in a dose-dependent manner (Fig. 1, D and E). In contrast, HCQ had no influence on the thermal stability of SARS-CoV-2 M^pro^ (Fig. 1, D and F). DSF is a powerful tool in early drug discovery (22) with the basic principle that drugs which bind to the therapeutic protein target will stabilize it and cause a positive shift in its Tm. However, small-molecule inhibitors have been shown to cause negative shifts in the Tm values of target proteins by disrupting their oligomeric interfaces, leading to thermal destabilization and subsequent loss of interaction between the protein subunits (23,24). Some examples include 6-hydroxy-DL-dopa binding to RAD52 (25) and SPD304 binding to TNF-α (26). In other cases, a ligand can bind more strongly to the non-native state than the native state of its target protein, such as that of Zn^2+^ and porcine growth hormone (27). To test whether TFQ binding disrupts the dimerization interface or binds to the non-native state of SARS-CoV-2 M^pro^, various biophysical methods were utilized to assess the conformational changes of SARS-CoV-2 M^pro^. Analytical ultracentrifugation studies revealed identical sedimentation coefficient at various concentrations of TFQ, suggesting the absence of dimer-to-monomer conversion of SARS-CoV-2 M^pro^ in the presence TFQ (Fig. 2A). Interestingly, results from circular dichroism (CD) spectroscopy revealed an increase in the far-UV signals (molar ellipticity at 222 nm) with increasing concentrations of TFQ, indicating that the total helical content of SARS-CoV-2 M^pro^ decreased upon TFQ binding (Fig. 2B). The decreased α-helical content was accompanied by reduced M^pro^ protease activity (Fig. 2, B and C). Together, these data suggested that TFQ may cause a local conformational change within its binding site, disrupting nearby α-helices and subsequently reducing M^pro^’s protease activity (Fig. 2, B and C). Moreover, because the sedimentation coefficient remained unchanged with TFQ, it is unlikely that TFQ caused unfolding of the overall structure of SARS-CoV-2 M^pro^ (Fig. 2A). To confirm this assumption, the solubility and stability of SARS-CoV-2 M^pro^ in the presence of different concentrations of TFQ or HCQ were tested. Consistently, the protein band of SARS-CoV-2 M^pro^ remain intact at concentration of TFQ up to 90 μM (Fig. 2D). At concentration above 120 μM TFQ, the soluble fraction of SARS-CoV-2 M^pro^ diminished gradually, suggesting that the conformational change induced by TFQ may expose some hydrophobic residues and ultimately result in protein aggregation (Fig. 2D). By contrast, HCQ do not influence the stability of SARS-CoV-2 M^pro^ at concentration up to 500 μM (Fig. 2D).

**Fig. 2.**
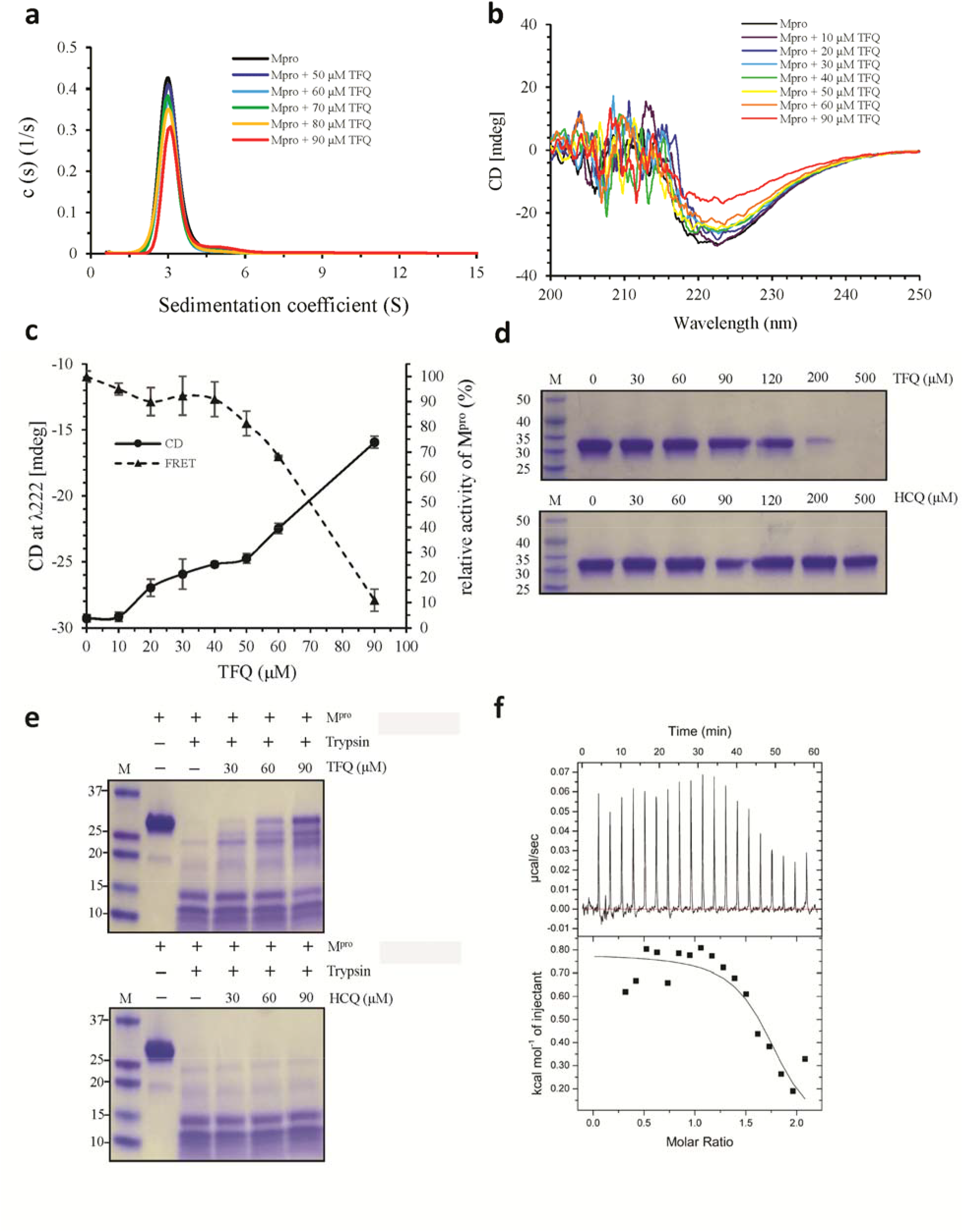
Deciphering the inhibition mechanism of TFQ on SARS-CoV-2 M^pro^. (**A**) Analytical ultracentrifugation (AUC) experiment of SARS-CoV-2 M^pro^ in the presence of different concentrations of TFQ. (**B**) The circular dichroism spectra of SARS-CoV-2 M^pro^ in the presence of different concentrations of TFQ. (**C**) Comparison of the far-UV CD signals (molar ellipticity at 222 nm) with enzyme activity of SARS-CoV-2 M^pro^ from FRET-base assay with increasing amounts of TFQ. The results are shown as a solid line (CD signals) or dashed line (enzyme activity measured by FRET) with error bars from at least two replicates. (**D**) SDS-PAGE detection of soluble fractions of SARS-CoV-2 M^pro^ in the presence of different concentration of TFQ or HCQ. (**E**) Limited proteolysis of SARS-CoV-2 M^pro^ by trypsin in the presence of different concentrations of TFQ (top) or HCQ (bottom). (**F**) Isothermal titration calorimeter (ITC) analysis of TFQ binding to SARS-CoV-2 M^pro^.

To further probe the conformational changes of SARS-CoV-2 M^pro^, we performed a limited proteolysis assay by trypsin digestion (28). The cleavage pattern indicated a greater degree of protection of SARS-CoV-2 M^pro^ from trypsin digestion at higher concentrations of TFQ (Fig. 2E). In contrast, no concentrations of HCQ tested reduced the cleavage of SARS-CoV-2 M^pro^ by trypsin digestion (Fig. 2E). Results from binding constant measurement by isothermal titration calorimetry (ITC) indicated TFQ bound to SARS-CoV-2 M^pro^ with micromolar affinity (*Kd* = ~10^−5^ M, Fig. 2F). These findings further supported the notion that TFQ binding induces local conformational changes in M^pro^ that trigger an active-to-inactive form transition, reduce its Tm and protease activity, and render it more resistant to trypsin digestion.

### Binding mode of TFQ to SARS-CoV-2 M^pro^

To elucidate the inhibitory mechanism of TFQ against SARS-CoV-2 M^pro^, molecular docking was performed using SwissDock (29). The resulting complex showed that TFQ fits well in the substrate-binding site of SARS-CoV-2 M^pro^ (Fig. 3A) and forms three polar contacts with F140, E166, and the active-site residue C145 (Fig. 3, A and C). In addition, eight residues of SARS-CoV-2 M^pro^, including H41, M49, L141, N142, S144, H164, M165 and Q189, contribute to the hydrophobic interface enclosing TFQ in the substrate-binding site (Fig. 3, A and C). To compare the differences between TFQ and HCQ at the molecular level, we also conducted a docking analysis of HCQ to SARS-CoV-2 M^pro^. Compared with TFQ, HCQ forms only one polar contact with E166 and fewer hydrophobic interactions within the substrate-binding site of SARS-CoV-2 M^pro^ (Fig. 3, B and D). The enhanced binding affinity of TFQ to SARS-CoV-2 M^pro^ may be attributed to the binding of the 3-(trifluoromethyl) phenoxy moiety of TFQ in the S1′ site of M^pro^, which is absent in the HCQ M^pro^ docked complex. Recently, the crystal structure of SARS-CoV-2 M^pro^ in complex with a mechanism-based inhibitor N3, was published (30). We compared the N3 inhibitor-M^pro^ complex (PDB code: 6LU7) with the TFQ-M^pro^ docked complex and showed that TFQ occupies the sites equivalent to P2, P1 and P1′ of the N3 inhibitor in the substrate-binding site (Fig. 3E). The S1 subsite of M^pro^ is highly specific to Gln at the P1 position of peptide substrate. The pentan-1,4-diamine moiety of TFQ appears to meet this requirement by mimicking the Gln side chain to form two hydrogen bonds with the side chain of E166 and main chain of F140 (Fig. 3, C and E). The hydrophobic quinoline core of TFQ is in close proximity to the S2 subsite, which prefers hydrophobic residues (Fig. 3E). In conclusion, the docking studies showed that TFQ binds to SARS-CoV-2 M^pro^ by mimicking its preferred peptide substrate.

**Fig. 3.**
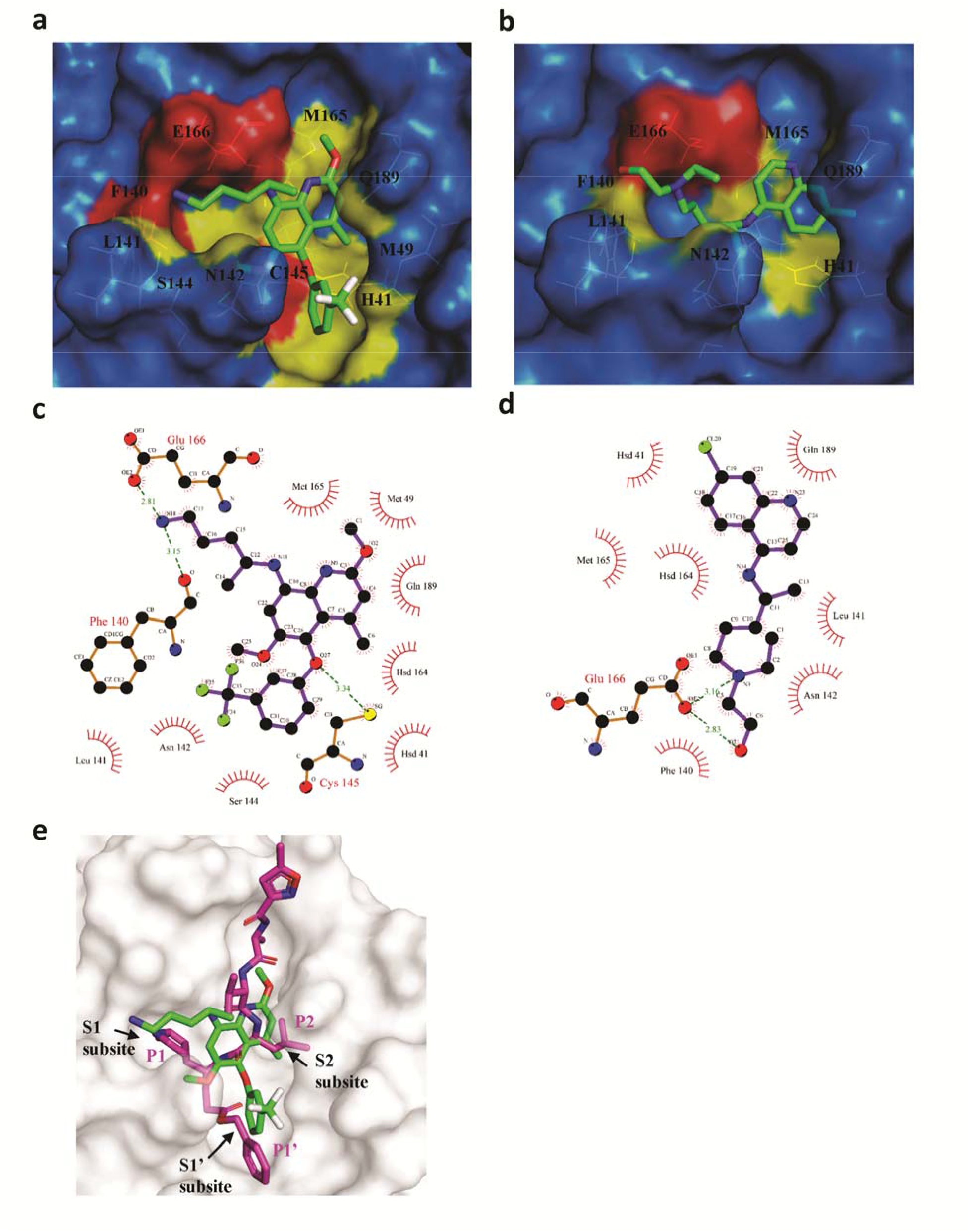
Molecular docking of TFQ or HCQ to SARS-CoV-2 M^pro^. Surface presentation of the substrate-binding pocket of SARS-CoV-2 M^pro^ bound with TFQ (**A**) or HCQ (**B**). Residues participated in H-bond formation are shown in red and hydrophobic interactions in yellow. Compounds are shown with ball-and-stick model with carbon atom in green. Detailed view of the interaction between TFQ (**C**) or HCQ (**D**) with SARS-CoV-2 M^pro^. (**E**) Superposition of the crystal structure of M^pro^-N3 (PDB: 6LU7) with M^pro^-TFQ docked complex. N3 inhibitor is shown with ball-and-stick model with carbon atom in magenta. The positions, P, on the peptide substrate, are named as P1, P2 (N-terminal to the cleavage site) and P1′ (C-terminal to the cleavage site) and the corresponding binding subsites located on the substrate-binding pocket are named as S1 (for binding to P1), S2 (binding to P2) and S1′ (for binding to P1′).

### TFQ inhibits SARS-CoV-2 production in cell culture system

Since TFQ treatment inactivates SARS-CoV-2 M^pro^, a key protease for viral replication in host cells, we examined the antiviral efficacy of TFQ on the viral production and infection rates of SARS-CoV-2. Vero E6 cells, which are kidney epithelial cells isolated from an African green monkey and commonly used to produce SARS-CoV-2 stocks in many research groups (16,31), were infected with SARS-CoV-2 (strain NTU02, GenBank:MT066176.1) at a multiplicity of infection (MOI) of 0.001 in the presence of TFQ (2.5 μM and 5 μM) or DMSO (control). Cells were subjected to two modes of drug treatment, one in which cells were pre-treated with TFQ for an hour prior to viral infection (full-time treatment), and the other in which cells were treated with TFQ after virus infection (post treatment) (Fig. 4A). After infection and TFQ treatment, cell supernatants were collected for further quantification of virus yield on Day 1, Day 2, and Day 3 after infection. The inhibition rate of TFQ against SARS-CoV-2 was determined by measuring viral RNA of nucleoprotein (N) using quantitative real-time RT-PCR (qRT-PCR). The results indicated that TFQ significantly repressed the yield of viral RNA in cell supernatant on day 1 to 2 after infection (Fig. 4B). Regardless of the treatment method used, the inhibition rate against viral RNA production was approximately 0–3.5% and 51.9–54% with 5 μM and 2.5 μM TFQ, respectively, at 48-hour post infection, implying the half maximal effective concentration (EC_50_) of TFQ was around 2.5 μM (Fig. 4C). Viral infection can lead to changes in cell morphology and death of host cells, also known as cytopathic effect (CPE) (32,33). Vero E6 cells are susceptible to SARS-CoV-2 infection, which induces CPE (34). We observed a significant decrease in SARS-CoV2-induced CPE in Vero E6 cells treated with 5 μM TFQ treatment compared with the DMSO treatment group (Fig. 4D), indicating that TFQ mitigates cell damages caused by SARS-CoV-2. Therefore, the data in Figure 4b observed on day 3 showed that there is no significant difference of viral RNA between DMSO-treated and TFQ (2.5 μM)-treated groups because the former lacked sufficient number of surviving host cells for virus production. Collectively, these data demonstrated that TFQ potently reduces SARS-CoV-2 production in the host cells.

**Fig. 4.**
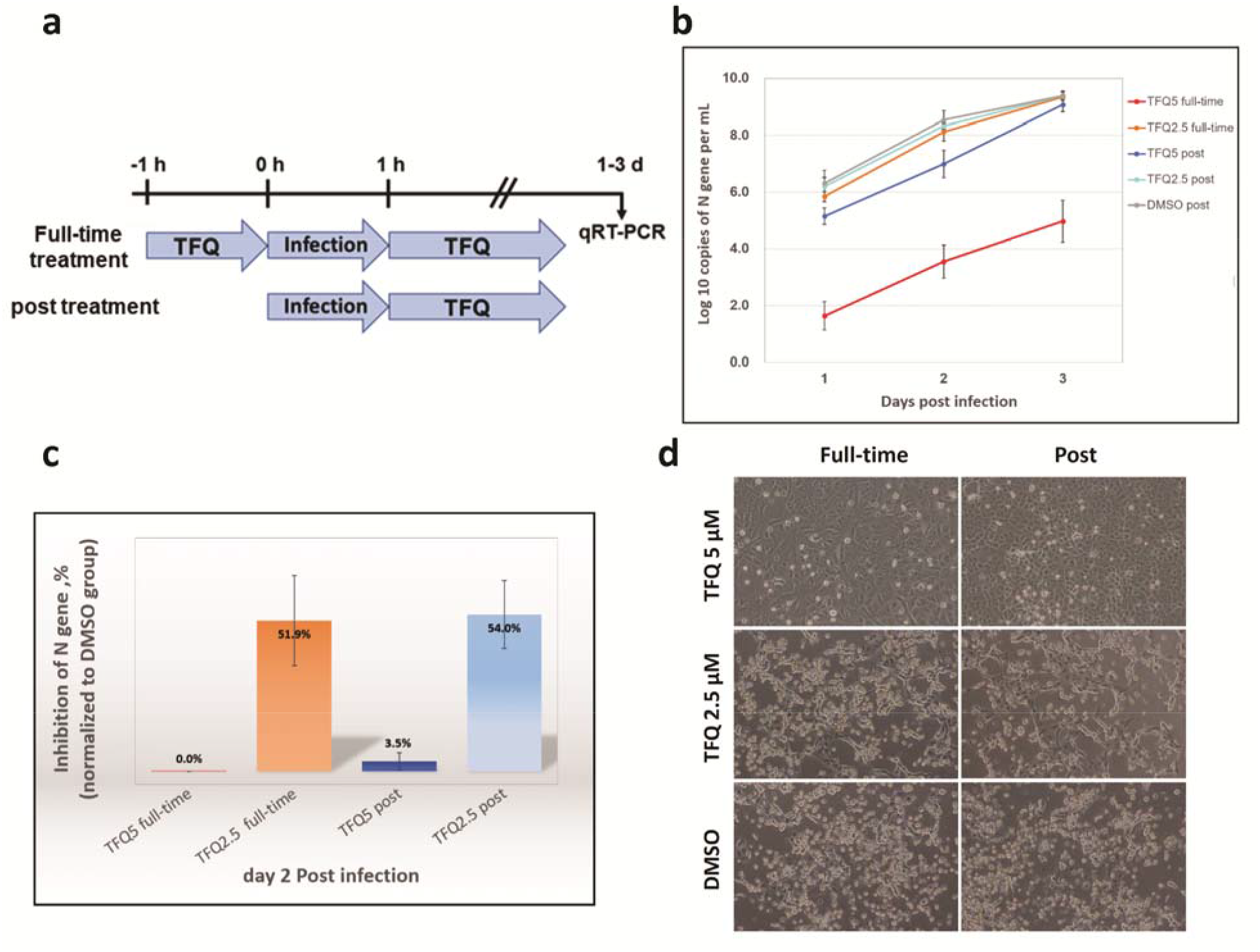
TFQ represses SARS-CoV-2 infection in Vero E6 cells. (**A**) A schematic illustrating two methods of treatment of Vero E6 cells infected SARS-CoV-2 with TFQ. In the pre-treatment group, cells were treated TFQ for one hour prior to viral infection. (**B**) The virus-infected Vero E6 cells were treated with TFQ (2.5 and 5 μM) or DMSO. The cell supernatant was collected on day 1, day 2, and day 3 and then subjected to qRT-PCR to determine the viral titer (n = 3). Data are shown as mean ± SD. (**C**) The inhibition rate of virus infection on day 2 in Vero E6 cells treated with 2.5 μM or 5 μM TFQ with full-time or post treatment (n = 3). Data are shown as represent mean ± SD. (**D**) 10× phase contrast images of virus-infected Vero E6 treated with DMSO or TFQ (2.5 μM and 5 μM) with TFQ full-time or post treatment.

## Discussion

Drug repurposing is an efficient way to accelerate the development of therapies for COVID-19. Here, we identified TFQ as a potent drug that inhibits SARS-CoV-2 replication by targeting its M^pro^ from an FDA-approved compound library. We first demonstrated that TFQ inhibits the enzymatic activity of SARS-CoV-2 M^pro^ by using a FRET-based assay. Subsequent molecular docking study indicated that TFQ binds directly to the substrate-binding pocket of SARS-CoV-2 M^pro^ as a competitive inhibitor. Moreover, binding of TFQ prevented M^pro^ from trypsin degradation and induced a negative shift in its Tm, supporting the conversion of SARS-CoV-2 M^pro^ from an active to inactive form in the presence of TFQ. Using CD spectroscopy, we showed that increasing TFQ concentrations reduced the α-helical content of M^pro^, suggesting possible unraveling of some α-helices. However, the results from trypsin digestion indicated that TFQ binding rendered M^pro^ more resistant to trypsin digestion, which indicates the presence of a more ordered structure, preventing it from trypsin digestion. In addition, the results from analytical ultracentrifugation (Fig. 2A) also suggested the formation of an ordered structure of the TFQ-M^pro^ complex. Thus, inhibition of M^pro^ by TFQ seems to convert M^pro^ from an active to inactive conformation. This mechanism differs from the typical mechanism of action of inhibitors that bind to the active site of the enzyme to block substrate binding. Therefore, we proposed a model shown in Fig. 5, illustrating the possible inhibition mechanism of TFQ on SARS-CoV-2 M^pro^. Based on the results shown in this study, the binding of low concentration (10 to 90 μM) of TFQ to M^pro^ decreases its protease activity by reducing the α-helical structure content. The conformational change may destabilize SARS-CoV-2 M^pro^ by exposing some hydrophobic residues to solvent, resulting decreased thermal stability. However, at concentration above 120 μM TFQ, the exposed hydrophobic region of SARS-CoV-2 M^pro^ may exceed a threshold, leading to protein aggregation and precipitation. Therefore, TFQ may inhibit the function of M^pro^ through a two-step progressive process to reduce SARS-CoV-2 production (Fig. 5).

**Fig. 5.**
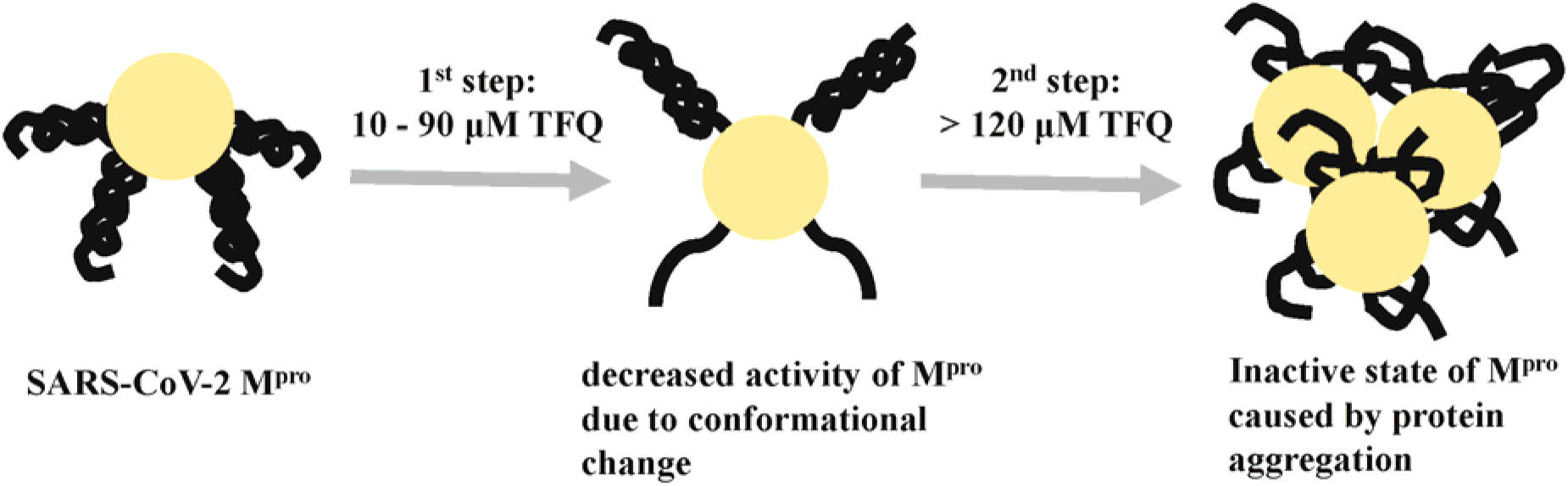
Model of TFQ-induced conformational changes in SARS-CoV-2 M^pro^. SARS-CoV-2 M^pro^ undergoes a local conformational change with increasing concentrations of TFQ (10 to 90 μM). Binding of TFQ alters nearby α-helices and to some extent reduces the proteolytic activity of SARS-CoV-2 M^pro^. At concentration above 120 μM TFQ, the exposed hydrophobic region of SARS-CoV-2 M^pro^ exceed a threshold, causing protein aggregation.

It is interesting to note that the N3 inhibitor blocked SARS-CoV-2 at a concentration of 10 μM in cell-based assay (30) whereas TFQ exhibited strong antiviral effect at a concentration of 5 μM in SARS-CoV-2 infected Vero E6 cells (Fig. 4). In contrast to TFQ which can be immediately evaluated in patients with COVID-19 in clinical trials, there is currently no safety, oral bioavailability, or pharmacokinetics study of N3 inhibitor in patients.

TFQ is approved for prophylaxis and treatment of malaria in the U.S. and Australia (15,35). As a preventive measure, a dose of 200 mg TFQ is recommended for three days prior to traveling and 200 mg per week until one week after return. For radical cure, a single dose of 300 mg TFQ is recommended (https://www.nc.cdc.gov/travel/news-announcements/tafenoquine-malaria-prophylaxis-and-treatment). Those above dose recommendations for malaria prophylaxis suggested that TFQ at higher doses may be tolerated by the human body. Contrary to the long half-life (one month or longer) and possible severe side effects, such as bulls-eye maculopathy, dry eye, nausea, diarrhea, anemia, liver failure, and muscle paralysis, of CQ and HCQ, the half-life of TFQ is relatively shorter (~14 days) and the side effects are less severe (36,37). Together with our data showing that 5 μM TFQ strongly inhibited SARS-CoV-2 infection *in vitro* (Fig. 4), especially when applied as TFQ pre-treatment (full-time treatment) to mimic the prophylactic use against viral infection, the repurposing of TFQ for the prevention and treatment of COVID-19 is worth looking into for clinical evaluations.

Although FDA-approved drugs that target M^pro^ have been identified using virtual docking methods (38,39), they have not been evaluated for their effects on M^pro^ protease activity by functional assays. It is worth mentioning that among those docking-positive candidates, none of them that we tested (at least 10; Table 1) showed strong inhibitory effects against the M^pro^ protease activity. Hence, the current study not only identifies the first anti-SARS-CoV-2 M^pro^ drug that has been tested functionally to inhibit M^pro^ protease activity and evaluated for safety (FDA-approved drug) but also provides a mechanistic insight into the mode of action of TFQ against SARS-CoV-2 M^pro^. The low clinical toxicity and mechanism-driven antiviral activity of TFQ against SARS-CoV-2 should warrant further testing in clinical trials.

**Table 1.** List of docking-positive FDA-approved compounds against SARS-CoV-2 M^pro^.

| Name | PMIDs |
| --- | --- |
| Lopinavir | 32280433 |
| Ritonavir | 32280433 |
| Atazanavir | 32280433 |
| Indinavir | 32280433 |
| Nelfinavir | 32296570 |
| Darunavir | 32280433 |
| Simeprevir | 32280433 |
| Saquinavir | 32280433 |
| Colistin | 32296570 |

## Materials and Methods

### Recombinant protein preparation

The full-length gene encoding SARS-CoV-2 main protease (M^pro^, ORF1ab polyprotein residues 3264-3569, GenBank code: MN908947.3) with *Escherichia coli* codon usage was synthesized and subcloned into pSol SUMO vector using Expresso® Solubility and Expression Screening System (Lucigen). A pET16b plasmid encoding the fluorescent protein substrate of M^pro^ (His_10_-mTurquoise2-TSAVLQSGFRKM-mVenus) was synthesized and constructed for FRET based high-throughput screening assay. Each expression plasmid was transformed into *E. coli* BL21 (DE3) and then grown in Luria Broth medium at 37 °C until OD_600_ reached between 0.6 and 0.8. Overexpression of M^pro^ or its fluorescent protein substrate was induced by the addition of 20% L-rhamnose or 0.5 mM IPTG and incubated for 18 hours at 20 °C. The cell pellets were resuspended in sonication buffer [50 mM Tris-HCl pH 8.0, 500 mM NaCl, 10 % glycerol, 1 mM tris(2-carboxyethyl)phosphine (TCEP), 1 mM phenylmethylsulfonyl fluoride (PMSF)] and lysed by sonication on ice. Following centrifugation at 28,000 g, 4 °C for 30 min, the supernatant was loaded onto a HisTrap FF column (GE Healthcare), washed by sonication buffer containing 10 mM imidazole, and eluted with a 20–200 mM imidazole gradient in sonication buffer. An adequate amount of TEV protease was added to remove the N-terminal SUMO fusion tag of M^pro^. Both TEV protease and His_6_-SUMO fusion tag were then removed by HisTrap FF column. The M^pro^ and its substrate protein were further purified by size-exclusion chromatography and stored in buffer containing 50 mM Tris-HCl pH 8.0, 200 mM NaCl, 5 % glycerol, and 1 mM TCEP. For solubility and stability test, SARS-CoV-2 M^pro^ was incubated with different concentration of TFQ or HCQ (30-500 μM) at room temperature for 30 mins. After centrifugation at 16000g for 3 mins, 20 ul of the supernatant of was boiled at 95℃ for 10 mins, and analyzed by 10 % SDS-PAGE (Bio-rad).

### Small-molecule compound library

Three small-molecule compound libraries, including the FDA-approved Drug Library, Clinical Compound Library, and Anti-COVID-19 Compound Library (MedChemExpress), were used to screen for drugs against SARS-CoV-2 M^pro^.

### Fluorescence resonance energy transfer (FRET) assay

SARS-CoV-2 M^pro^ (4 μM) in assay buffer (20 mM Tris-HCl 7.8, 20 mM NaCl) was pre-incubated with or without 60 μM compounds for 30 min at room temperature in 96-well black Optiplate. The reaction was initiated by addition of 20 μM fluorescent protein substrate. Substrate cleavage was monitored continuously for 1 hour by detecting mTurquoise2 fluorescence (excitation: 434 nm / emission: 474 nm) using Synergy™ H1 hybrid multi-mode microplate reader (BioTek Instruments, Inc.). The first 15 min of the reaction was used to calculate initial velocity (V_0_) by linear regression. The calculated initial velocity with each compound was normalized to DMSO control. The IC_50_ was calculated by plotting the initial velocity against various concentrations of TFQ by use of a dose-response curve in Prism 8 software.

### Protein thermal shift assay using differential scanning fluorimetry

Differential scanning fluorimetry (DSF) assays were conducted as previously described (Lo, et al., 2004). Briefly, the experiment was carried out on a CFX96 RT-PCR instrument (Bio-Rad) in a buffer comprising 25 mM Tris pH 8.0, 150 mM NaCl, 5X SYPRO Orange dye (Sigma-Aldrich), and 8 μM SARS-CoV-2 M^pro^ in the presence or absence of 120 μM compound in each well. Fluorescence was monitored when temperature was gradually raised from 25 to 90 °C in 0.3 °C increments at 12-second intervals. Melt curve data were plotted using the Boltzmann model to obtain the temperature midpoint of unfolding of the protein using Prism 8.0 software (GraphPad).

### Circular dichroism (CD) spectroscopy

CD signals were measured using a Jasco J-815 spectropolarimeter with 0.1-cm quartz cuvettes and a 1-mm slit width. The molar ellipticity at 222 nm of all samples was recorded to analyze the protein conformational changes at different concentrations of TFQ (10–500 μM). All spectra were corrected for buffer absorption.

### Analytical ultracentrifugation (AUC)

To assess the quaternary structural changes of SARS-CoV-2 M^pro^ in the presence of TFQ, sedimentation velocity experiments were performed using a Beckman Optima XL-A analytical ultracentrifuge (Beckman Coulter, CA, USA). Before ultracentrifugation, the protein sample was preincubated with various concentrations of TFQ (50–100 μM) at room temperature for 30 min. The protein sample and buffer solutions (25 mM Tris 8.0, 150 mM NaCl) were separately loaded onto the double sector centerpiece and placed in a Beckman An-50 Ti rotor. The experiments were performed at 20 °C and at a rotor speed of 42,000 rpm. The protein samples were monitored by the UV absorbance at 280 nm in continuous mode with a time interval of 480 s and a step size of 0.002 cm. Multiple scans at different time points were fitted to a continuous size distribution model by the program SEDFIT (Schuck et al., 2002). All size distributions were solved at a confidence level of p = 0.95, a best fitted average anhydrous frictional ratio (f/f0), and a resolution N of 250 sedimentation coefficients between 0.1 and 20.0 S.

### Limited proteolysis by trypsin

The protein sample was preincubated with various concentrations of TFQ (30–120 μM) at room temperature for 30 min. Proteolysis was then performed by mixing SARS-CoV-2 M^pro^ (0.8 mg/ml) with trypsin at a protease-to-protein ratio of 1:10 (w/w) in reaction buffer (25 mM Tris 8.0, 150 mM NaCl) at 37 °C for 30 min. The reaction was stopped by adding SDS sample loading buffer and boiling at 95 °C for 10 min and subjected to SDS-PAGE (4–20%).

### Isothermal titration calorimetry

The binding of TFQ to SARS-CoV-2 M^pro^ were conducted on an ITC-200 instrument (MicroCal, Northampton, MA, USA) at 25 °C. SARS-CoV-2 M^pro^ and the inhibitors were dissolved in the assay buffer (20 mM Tris pH 8.0, 20 mM NaCl, 0.5 % DMSO). Two microliters aliquots of TFQ at concentration of 500 μM in the syringe were injected into the cell containing 50 μM M^pro^ at 3-min intervals. Data were fit to a one-site binding model using the commercial Origin 7.0 program to obtain the thermodynamic parameters.

### Molecular docking

Molecular docking of TFQ and HCQ to SARS-CoV-2 M^pro^ was performed using SwissDock and the crystal structure of SARS-CoV-2 M^pro^ (PDB code: 6LU7). The docking pose with the highest docking score for each compound was selected for further analysis. Ligand plot of each compound was generated by PDBsum.

### Virus and cell culture

SARS-CoV2 (strain NTU02, GenBank:MT066176.1) was isolated from a COVID-19 patient at National Taiwan University Hospital and grown in Vero E6 cells. Cells were maintained in Dulbecco’s modified Eagle’s medium (DMEM) containing 10% fetal bovine serum (FBS).

### Viral infections

Vero E6 cells (1 × 10^7^) were washed with PBS, incubated with virus diluted in serum-free DMEM containing tosylsulfonyl phenylalanyl chloromethyl ketone (TPCK)-trypsin (2 μg/ml) for 1 hour at 37 °C at a multiplicities of infection (MOI) of 0.001. One hour after infection, the virus inoculum was removed. The infected cells were washed with PBS and incubated with fresh medium containing 2% FBS.

### Time course assay of TFQ

Vero E6 (7 × 10^4^) cells were seeded in 24-well plates and subjected to two modes of drug treatment, one in which cells were pre-treated with drugs for an hour prior to viral infection, and the other without drug pre-treatment. Cells were then infected with virus for one hour in the absence of drugs. After infection, cells were washed with PBS, and cultured with drug-containing medium until the end of the experiment. The virus-containing supernatants were harvested at one to three days post-infection and subjected to qRT-PCR to determine the viral titers. The viral cytopathic effect (CPE) was observed under microscope and imaged at 3-day post infection.

### Viral RNA extraction and quantitative RT-PCR

The viral RNA in supernatant was extracted using the QIAamp Viral RNA Mini Kit (QIAGEN). The extracted RNA was reverse transcribed using SuperScriptTM III reverse transcriptase (Invitrogen). The cDNAs were amplified by real-time PCR using the LightCycler FastStart DNA Master HybProbe (Roche Molecular Biochemicals) with a Light Cycler® 96 (Roche Molecular Biochemicals) for 50 cycles of 10 sec at 95 °C, annealing of 10 sec at 58 °C, and elongation of 10 sec at 72 °C to detect N gene of SARS-CoV2. The following primers and probe were used:

N_Sarbeco_F1:5’-CACATTGGCACCCGCAATC-3’
N_Sarbeco_R1:5’-GAGGAACGAGAAGAGGCTTG-3’
N_Sarbeco_P1: 5’-FAM-ACTTCCTCAAGGAACAACATTGCCA-BHQ1-3’.

### Statistical analysis

Data of bar or curve graphs display as percentage or number compared to control groups with a standard deviation of two or three independent experiments. Microsoft Excel was used for statistical analyses. The two-tailed independent Student’s *t*-test was used to compare continuous variables between two groups. All experiments were carried out at least twice. The statistical significance level of all tests is set to 0.05.

## Supplementary Materials

### Supplementary Figures

Fig. S1. The purification of SARS-CoV-2 M^pro^.

Fig. S2. The purification of fluorescent protein substrate of SARS-CoV-2 M^pro^.

## Acknowledgments

We would like to thank Ms. Yi-Jen Liau at National Taiwan University for technical assistance of SARS-CoV-2 preparation and SCI Pharmtech Inc. for providing HCQ.

## Author contributions

Y.C. and W.-H.Y. designed and carried out the experiments, interpreted data, and wrote the manuscript; Y.-C.W., L.-M.H., C.-S.Y., Y.-L.L., M.-H.H., C.-L.T., Y.-Z.C., B.-Y.H., C.-F.H., Y.-L.H. and Y.-P.C. carried out the experiments; Y.-C.W., L.-M.H., C.-S.Y., Y.-L.L., J.-S.C., Y.-P.C., D.-Y.C., L.-B.J. and C.-H.T. analyzed and discussed the data; M.-C.H. supervised the entire project and prepared the manuscript.

## Data and materials availability

All data are presented in the paper or the Supplementary Materials. The materials used in this study should be requested form M.C.H.

